# Different outcomes of neonatal and adult Zika virus infection on startle reflex and prepulse inhibition in mice

**DOI:** 10.1101/2022.11.08.515594

**Authors:** Isis N. O. Souza, Brenda S. Andrade, Paula S. Frost, Romulo L. S. Neris, Daniel Gavino-Leopoldino, Andrea T. Da Poian, Iranaia Assunção-Miranda, Claudia P. Figueiredo, Julia R. Clarke, Gilda A. Neves

**Affiliations:** Laboratory of Molecular Pharmacology, Institute of Biomedical Sciences, Universidade Federal do Rio de Janeiro, Brazil; School of Pharmacy, Universidade Federal do Rio de Janeiro, Brazil; Institute of Medical Biochemistry Leopoldo de Meis (IBqM), Universidade Federal do Rio de Janeiro, Brazil; Institute of Microbiology Paulo de Goes, Universidade Federal do Rio de Janeiro, Brazil

**Author notes:** Corresponding author: Gilda A. Neves, Laboratory of Molecular Pharmacology, Institute of Biomedical Sciences, Federal University of Rio de Janeiro, Av. Carlos Chagas Filho, 373, bloco J, sala J1-029, CEP 21941-902, Cidade Universitária, Ilha do Fundão, Rio de Janeiro, RJ, Brazil.

**Keywords:** flavivirus, neurodevelopment, sensorimotor gating, neuromotor function, viral infection, arboviral, CNS viral diseases

## Abstract

Zika virus (ZIKV) infection causes severe neurological consequences in both gestationally-exposed infants and adults. Sensorial gating deficits strongly correlate to the motor, sensorial and cognitive impairments observed in ZIKV-infected patients. However, to date, no startle response or prepulse inhibition (PPI) assessment has been made in patients or animal models. In this study, we identified different outcomes for the age of infection and sex in wild-type mice: neonatally infected animals presented an increase in PPI and startle latency in both sexes, while adult infected males presented lower startle amplitude but preserved PPI. Our data further the understanding of the functional impacts of ZIKV on the developing and mature nervous system, which could help explain other behavioral and cognitive alterations caused by the virus. With this study, we support the use of startle reflex testing in ZIKV exposed patients, especially infants, allowing for early detection of functional neuromotor damage and early intervention.

## Introduction

The Zika virus (ZIKV) epidemic of 2015 was a public health emergency of international concern, infecting over 730 thousand people on the American continent (World Health Organization 2022). Severe neurological consequences were reported in adults, including acute encephalitis, Guillain-Barré syndrome, and long-term cognitive impairment (Mehta et al. 2018; Muñoz et al. 2017; Rozé, Najioullah, Signate, et al. 2016; Zucker et al. 2017). Moreover, the gestational infection had devastating effects on newborns, ranging from the well-publicized microcephaly to a myriad of anatomical, motor, and behavioral symptoms that make up the congenital ZIKV syndrome (Pereira et al. 2020; Moore et al. 2017). As those patients continue to grow, new symptoms and developments follow.

It is well known that viral infections, either in or out of critical developmental periods, can cause long-lasting effects on brain circuitry and function (de Vries 2019; van den Pol 2006). For example, cohorts of ZIKV-infected babies, with and without microcephaly, have been monitored by numerous research groups. These studies revealed a consistent failure to reach developmental landmarks in social, communication, and cognitive skills (Alves et al. 2018; Mulkey et al. 2020), as well as a high incidence of motor impairments and drug-resistant epileptic seizures (A. L. de Carvalho et al. 2020; M. Carvalho et al. 2020). Moreover, several case reports of ZIKV-induced encephalitis in adults include consciousness alterations and cognitive impairments that may last for weeks (Rozé, Najioullah, Fergé, et al. 2016; Mehta et al. 2018; Zucker et al. 2017). Thus, studies trying to understand better the neural mechanisms underlying these changes are still needed.

Sensorimotor gating is a key mechanism for filtering sensory information in order to maintain adequate attentional processes for daily life activities (Braff and Geyer 1988). Impairment in sensorimotor gating is present in several neurological and neuropsychiatric disorders (Geyer 2006), including those of viral etiology (Asp et al., 2010; Minassian et al. 2013; Liu et al., 2014; Ronca, Dineley, and Paessler 2016). Sensorial gating deficits strongly correlate to motor, sensorial and cognitive impairments, so they can either explain or be caused by alterations in these domains (Geyer 2006). The prepulse inhibition (PPI) paradigm, in which a weak sensory stimulus presented before a stronger salient one reduces the motor reflex, is a robust and conserved method to evaluate sensorimotor gating in mammals. However, no startle response or PPI assessment was made in adults or newborn ZIKV-infected patients, nor was this feature investigated in animal models of ZIKV infection.

Previous papers from our group developed useful animal models to study ZIKV infection outcomes in mice. Our works showed that infection with a Brazilian strain of ZIKV causes persistent neurological effects in the neonatal and adult mouse, including long-lasting motor and cognitive impairment (Souza et al. 2018; Figueiredo et al. 2019; Pinheiro et al. 2020). Furthermore, we identified synaptic damage to the hippocampus (Figueiredo et al. 2019) as well as long-lasting lesions in the striatum, hippocampus, thalamus, and cerebral cortex (Souza et al. 2018), all brain regions involved in PPI modulation (Gómez-Nieto, Hormigo, and López 2020). Therefore, we hypothesized that ZIKV infection could impair startle response and PPI. In this work, we investigated whether infection with a Brazilian strain of ZIKV in either the mature or developing nervous system could cause long-term modifications in those measures, furthering our understanding of ZIKV infection and opening paths for early intervention.

## Methods

### Animals

Swiss mice of both sexes were bred and kept at the Center of Health Sciences, Federal University of Rio de Janeiro animal facilities. For neonatal infections, three days old (PND3) litters (5 for each condition) were standardized to eight to ten pups each and, whenever possible, were composed of equal numbers of males and females. Litters were allocated to experimental groups by simple randomization. Infections were performed as described below. At 21 days of age, animals were weaned and housed with mice of the same litter and sex, with two to five animals per cage. For adult infections, 2.5-3 months old mice were used. All animals were housed in polypropylene cages containing pine shavings as bedding, kept at 23 ± 2°C, in a twelve-hour light-dark cycle (lights on 6 am), and had unrestricted access to filtered water and a standard diet. All experiments were performed according to the U.S. National Research Council’s Guide for the Care and Use of Laboratory Animals and the Brazilian National Council for Animal Experimentation Control (CONCEA) guidelines. The local ethics committee previously approved all animal procedures (CEUA/UFRJ protocols number 043/2016, 052/2017, and 126/2018).

### Virus

ZIKV isolated from a febrile patient in the state of Pernambuco, Brazil (BRPE243/2015, KX197192) was propagated in the C6/36 cell lineage cultured in Leibovitz L-15 medium (Invitrogen) supplemented with 5% fetal bovine serum (FBS) at 28°C, as previously described in Coelho et al. (Coelho et al. 2017) and stored at −80°C. The titer of viral stock was determined by plaque assay in Vero cells cultured in high-glucose Dulbecco modified Eagle medium (DMEM; Invitrogen) after 10-fold serial dilutions. The same volume of virus-free conditioned medium of C6/36 cells cultured in the same conditions was used in control groups (Mock).

### Infections

For neonatal infections, pups at PND3 were injected subcutaneously with 30 μL of ZIKV (10^6^ plaque-forming units) or Mock medium at the same dilution. The viral titer was chosen according to our previous work (Souza et al. 2018). All pups of the same litter received the same intervention to avoid cross-contamination. Animals with signs of hemorrhage or reflux were excluded from experimentation. For adult infections, animals were anesthetized with 2.5% isoflurane (Cristália; São Paulo, Brazil) using a vaporizer system (Norwell, MA, USA) and were gently manually restrained only during the injection procedure. To access the lateral ventricle, a 2.5 mm-long needle was inserted unilaterally 1 mm to the right of the midline point equidistant from each eye and parallel to a line drawn through the anterior base of the eye (Figueiredo et al. 2019). Next, three microliters of ZIKV (10^5^ plaque-forming units) or Mock medium were slowly infused using a Hamilton syringe. Mice showing any signs of misplaced injections or brain hemorrhage were excluded from experimentation.

### Prepulse inhibition of startle reflex

As previously described (Marques et al. 2020), the test was performed in a soundproof chamber containing a sound generator and an accelerometer for motion detection (Panlab, Harvard Instruments). Neonatally-infected animals were evaluated at ∼90 dpi, while adult-infected animals were evaluated at 1, 6, 14, and 45 dpi. Background noise at 65 dB was played throughout the experiment. Initially, the animals were habituated to background noise for 5 min, and soon after, they were exposed to 5 pulses of white noise at 120 dB with 50 ms duration, a startle accommodation phase. These pulses were not included in the analyses. The experiment consisted of 5 stimulus blocks (repeated 10 times each) presented randomly at an interval of 20 ± 10 s. The blocks consisted of: no stimulus (background noise only), pulse (120 dB), and pulse preceded by prepulse at one of three intensities: 72, 80, or 90 dB (white noise, 20 ms long played 100 ms before pulse). The accelerometer trace was digitalized with Startle software (v.1.2.04, Panlab, Harvard Instruments), and the startle latency and amplitude were detected up to 500 ms after pulse presentation. The mean startle amplitudes of each block and the mean latency of pulse-only blocks were used for the analyses. Animals infected in the neonatal period were evaluated at PND95, while those infected in adulthood were evaluated 1, 6, 14, and 45 days post-infection (dpi).

### Statistical analysis

All data are expressed as mean ± standard error of the mean (S.E.M.). The percent inhibition of the startle reflex for each prepulse intensity was calculated using the equation: %PPI = 100 – (average startle from prepulse + pulse block/ average startle from pulse block × 100). Average PPI was calculated for each animal as the arithmetic mean of PPI in the three prepulse intensities. In the neonatal cohort, startle amplitude, startle latency, and average PPI data were analyzed using two-way ANOVA with group (Mock or ZIKV) and sex (male or female) as independent factors. Data from the adult cohort was analyzed by a three-way repeated measures ANOVA, with group and sex as independent factors and days post-infection (dpi) as the repetition factor. All ANOVAs were followed by Tukey *posthoc* test. Data stratified by prepulse intensities (Fig 1S and 2S), their statistical analysis, and the complete description of the ANOVAs results (Tables 1S-3S) are available in the supplementary material. Statistics were calculated using SPSS v. 17 (IBM), and data were plotted using Prism 8 (GraphPad). Differences were considered significant at *p* < 0.05. Experimenters were blind to experimental groups during data analysis.

## Results

### Neonatal *ZIKV infection delayed startle response and increased PPI*

ZIKV infection in the neonatal period, a critical stage of neurodevelopment, can cause long-lasting effects on brain circuitry (Stanelle-Bertram et al., 2018; Mavigner et al., 2018; Souza et al., 2018). Therefore, we hypothesized that neonatal ZIKV infection could cause late startle and PPI changes. We observed no significant differences in startle amplitude between Mock and ZIKV-infected groups, either in male or female mice (Figures 1a and 1d, *p* > 0.05). This observation indicates that, although motor impairments were found in that animal model (Souza et al. 2018), they did not impair the magnitude of the motor reflex caused by the auditory stimulus. Notwithstanding, even when startle amplitude is within normal range, there might be a delay in motor response identifiable as increased startle latency. Indeed, ZIKV-infected animals of both sexes showed a statistically significant increase in startle latency (Figures 1b and 1e, *p* = 0.007), which argues in favor of a change in the speed of neuronal processing. Surprisingly, the average PPI percentage was increased in ZIKV-infected mice compared to the control group (Figures 1c and 1f, *p* = 0.026). Finally, when accounting for individual prepulse intensities, a triple interaction was found to be significant (p = 0.026). Posthoc analysis showed a significant increase in PPI only in females at 80 dB (*p* = 0.003) and 90 dB (*p* = 0.004) prepulse (Sup Figure 1), showing that the increase in PPI observed has a most substantial contribution of female infected mice response.

**Figure 1.**
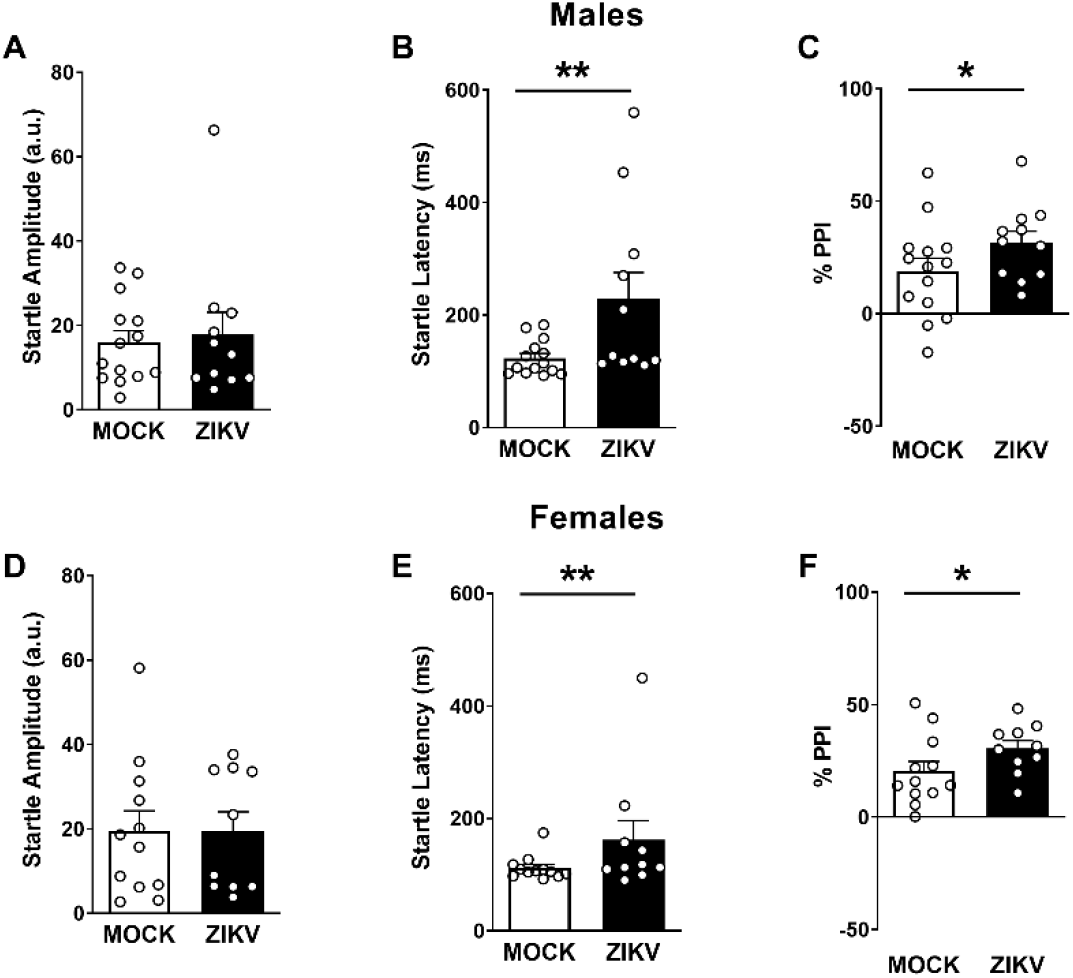
ZIKV infection in the developing brain causes alterations in startle latency and PPI. ZIKV did not change the startle amplitude of male **(A)** and female **(D)** mice (two-way ANOVA, *p* > 0.05) but increased the startle latency of both male **(B)** and female mice **(E)** (two-way ANOVA, F_group(1,43)_ = 8.149, *p* = 0.007). ZIKV increased the average percentage of prepulse inhibition (PPI) of both male **(C)** and female mice **(F)** (two-way ANOVA, F_group(1,43)_ = 5.315, *p* = 0.026). n = 14 Mock and 11 ZIKV for males and 12 Mock and 10 ZIKV for females. **p < 0.01, *p < 0.05 Mock vs. ZIKV (group effect). Data are expressed as mean ± S.E.M.

### Adult ZIKV infection impairs startle amplitude in male mice

ZIKV exhibits a preference for certain brain areas in our adult infection model, such as the hippocampus, the striatum, and the prefrontal cortex (Figueiredo et al. 2019), all coincidentally crucial for proper sensorimotor gating. Accordingly, we performed the same measurements as the neonatal model but evaluated several time points after infection. Evaluating startle amplitude, three-way ANOVA revealed a triple interaction effect (*p* = 0.042). Mock-injected males showed typical accommodation of startle amplitude over time (*p* < 0.001, *p* = 0.034, and *p* < 0.001 comparing 6, 14 and 45 dpi vs. 1 dpi, respectively). However, the same phenomenon was not detected in ZIKV-infected male mice (*p* = 0.440 for posthoc dpi comparisons), showing a significantly lower startle amplitude at 1 (p < 0.001) and 14 dpi (*p* = 0.008) in comparison to Mock-injected animals (Figure 2a). On the other hand, no startle accommodation or post hoc statistical differences were observed between female mice on none of the days evaluated (Figure 2d). Unlike the observation in the neonatal cohort, a significant sex difference in startle amplitude was detected in Mock-injected mice at 1 (*p* < 0.001) and 14 dpi (*p* = 0.019), but not in ZIKV-infected ones (*p* = 0.669).

**Figure 2.**
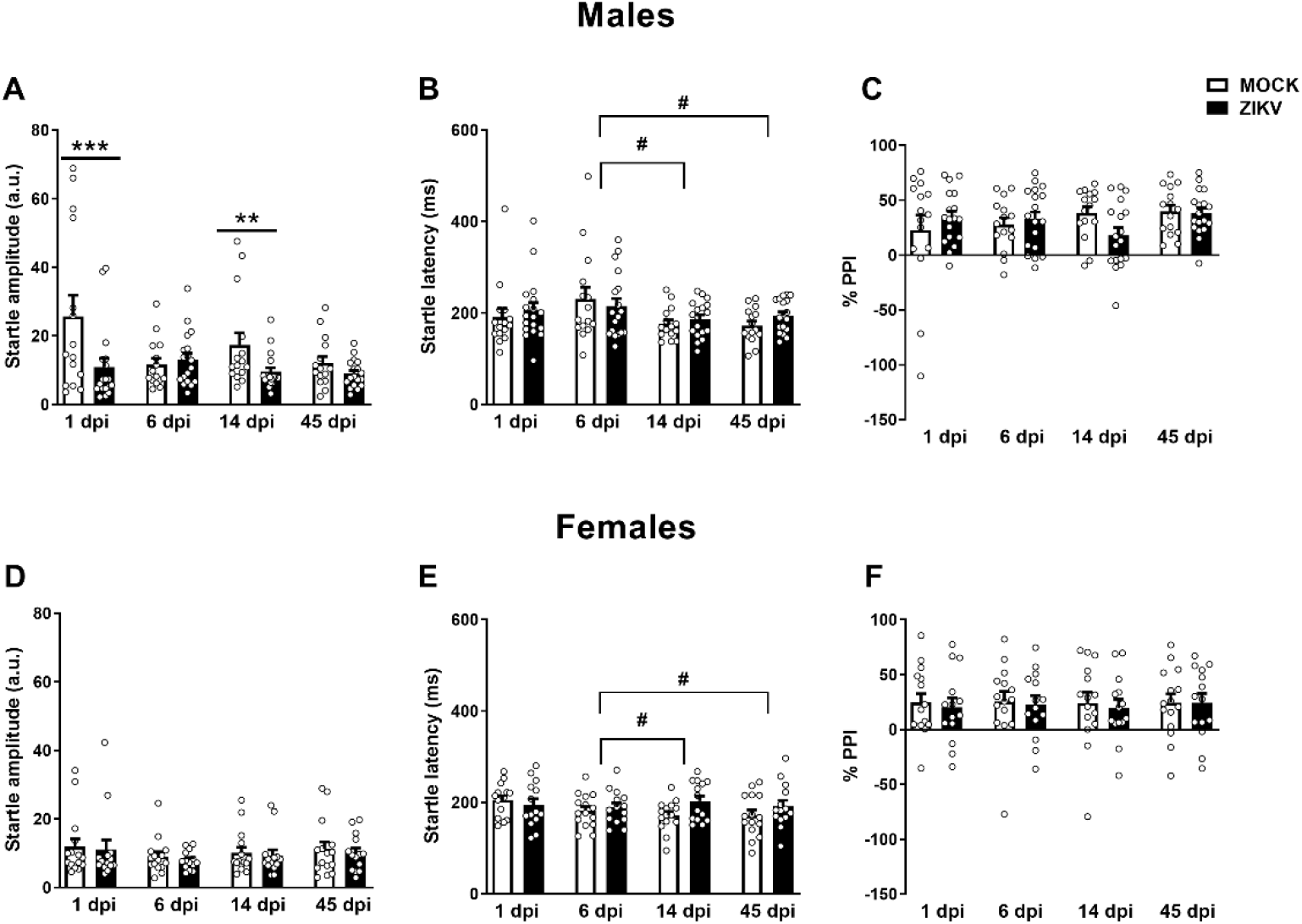
ZIKV infection in the mature brain causes alterations of startle parameters over time but preserves PPI. ZIKV reduced startle amplitude at 1 and 14 dpi in male **(A)** but not female **(D)** mice (three-way ANOVA, F_triple-interaction(3,174)_ = 2.785, *p* = 0.042). Startle latency was reduced in male **(B)** and female **(E)** mice over time, independent of the infection group (three-way ANOVA, F_dpi(3,174)_ = 2.845, *p* = 0.039). ZIKV did not alter the average percentage of prepulse inhibition (PPI) of either male **(C)** or female mice **(F)** (three-way ANOVA, *p* > 0.05). n = 15 Mock and 18 ZIKV for males and 15 Mock and 14 ZIKV for females. In A: \*\**p* < 0.01, ***p < 0.001 Mock vs. ZIKV, in B and E: ^#^p < 0.05 between dpi’s. Data are expressed as mean ± S.E.M.

In these same animals, we found a decrease in startle latency overtime, reaching statistical significance between 6 and 14 dpi (*p* = 0.041) and 6 and 45 dpi (*p* = 0.016) in both male and female mice (Figure 2b and 2e). This data shows an expected startle latency reduction already described after multiple evaluations in the same animals (Pilz and Schnitzler 1996). No differences were observed between groups (Mock vs. ZIKV) at any time point. Finally, we found no difference between groups in average PPI percentage across time points in either male or female mice (Figures 2c and 2f). When accounting for prepulse intensity in each timepoint, a significant increase in PPI was seen at the 72dB prepulse intensity between groups at 1 dpi only (*p* = 0.006) in a sex-independent manner (Sup Figure 2).

## Discussion

This work presents the first investigation of startle parameters and PPI in ZIKV-infection models. Our data showed different outcomes according to the neurodevelopmental stage when the infection occurs. While we observed no difference in startle amplitude in the neonatal infection model, there was a significant increase in startle latency in both male and female ZIKV-infected mice. The acute startle response is an important protective reflex against predators or impact (Koch 1999). It is mediated by a pontomedullary brainstem pathway that projects to the spinal cord, activating motor neurons (Gómez-Nieto, Hormigo, and López 2020). Startle latency changes are usually related to alterations in the speed of neuronal communication (Davis et al. 1982). Changes in motor nerve connectivity and conductivity were already described in models of systemic ZIKV infection (Morrey et al. 2019). Furthermore, sensorimotor maturation in rodents continues to develop along the first three postnatal weeks (Arakawa 2019). Therefore, considering the profound histological alterations caused by ZIKV infection at PND3 (Souza et al. 2018), it is possible to speculate that those areas and/or pathways were damaged during development and did not mature properly, increasing the time for neuronal processing and, therefore, the startle latency.

Surprisingly, though, we observed an increase in PPI in those animals. PPI “improvements” are rarely reported in the literature, being especially curious after a harmful intervention such as a viral infection, which often causes the opposite effect (Ozawa et al. 2006; Ronca, Dineley, and Paessler 2016). However, different genetic mutations have given rise to increases in PPI (Iscru et al. 2009; Kulikov et al. 2016). In these studies, an increase in startle amplitude was also observed; therefore, the PPI alteration was interpreted as a greater sensitivity to the prepulse. In our model, the startle response was similar in both groups, with no window for this interpretation. On the other hand, forebrain to brainstem pathways are capable of boosting attention to the prepulse, enhancing PPI independently of an increased pulse startle response. The areas involved include the lateral amygdala, the posterior parietal cortex, the primary auditory cortex, and the thalamus (Hazlett et al. 2001; Du, Wu, and Li 2011). Given the neuroanatomical and morphological changes in the brain of ZIKV-infected animals in both the cortex and thalamus (Souza et al. 2018), it is possible that adaptations in sensory and attentional information processing are involved in increasing PPI in our model. Evaluation of ZIKV-infected animals in attention tasks, as well as functional assessments in the brain areas involved in PPI modulation, can be of value in explaining those results.

In adult males, we observed a reduction in startle amplitude in some, but not all, days post-infection. In addition, a sex difference in the Mock-infected groups was also observed on these same days. These findings could be partially explained by the high intergroup variability in the Mock datasets at those time points. However, the transient decrease in startle response in ZIKV-infected male mice agrees with our previous data showing a motor coordination impairment. These changes might be related to the inflammatory process generated by the infection, including a lasting microglial activation (until 30 dpi) (Figueiredo et al. 2019). Furthermore, our data do not make clear whether the reduced startle amplitude observed in ZIKV-infected male mice is a consequence only of poor startle response or a combination with a lack of long-term startle habituation. An experiment specifically designed to investigate these parameters without prepulse exposure would help further elucidate this impairment.

Regarding startle latency, there was a significant reduction in both groups between 6, 14, and 45 dpi for both male and female mice. Habituation has also been described for startle latency when evaluated over time (Pilz and Schnitzler 1996), which argues that ZIKV infection has not disrupted that feature in adult animals. While average PPI was not altered in any time points for male and female animals, we observed a significant increase at 1 dpi with the 72 dB prepulse intensity when accounting for individual prepulse intensities. Again, intergroup variability in the Mock dataset seems to play a significant role in this finding.

There are, however, limitations to this study. While the acute startle response matures around PND12 (Koch 1999), we decided to evaluate neonatally infected animals only in adulthood, as our main goal was to identify the late consequences of the viral infection. Evaluation at different time points could help understand the chronology of the observed alterations. Moreover, we did not directly evaluate histological or cellular damage to the brainstem. Other groups with gestational and neonatal infection models have identified necrotic foci and cytoarchitectural alterations in the brainstem of ZIKV-infected animals (Snyder-Keller et al. 2019; Cugola et al. 2016; Paul et al. 2018), which corroborates our hypothesis. The used models also have some important features to be considered for data interpretation. In the neonatal model, direct infection in pups evades the effects of maternal immune activation. However, it still matches the timescale of several neurodevelopmental processes corresponding to humans’ third trimester of pregnancy (Semple et al. 2013; Clancy, Darlington, and Finlay 2001). Moreover, it also allows a more refined control of the viral titer exposure of each animal. In our adult mice model, injection is done intraventricularly, which allows better control of the viral titer that reaches the brain but also bypasses the peripheral immune response and reduces construct validity (Balint et al. 2021). On the other hand, peripheral infection in wild-type mice is often challenging due to increased interferon response, and using immunosuppressed animals defeats a peripheral infection’s purpose (Balint et al. 2021).

## Conclusion

By using the PPI paradigm, this study is the first to evaluate startle response and sensorimotor gating in ZIKV infection models. We identified different outcomes when accounting for age and sex, with neonatally infected animals presenting an increase in PPI and startle latency in both sexes, while adult infected males presented lower startle amplitude. Our data further the understanding of the functional impacts of ZIKV on the developing and mature nervous system, which might help explain other behavioral and cognitive alterations caused by the virus. With this study, we support the use of startle reflex testing in ZIKV-exposed patients, especially infants, allowing for early detection of functional neuromotor damage and early intervention.

## Abbreviations

ANOVA: analysis of variance
DMEM: Dulbecco modified Eagle medium
dpi: days post-infection
FBS: fetal bovine serum
PND: postnatal day
PPI: prepulse inhibition
S.E.M.: standard error of the mean
ZIKV: Zika virus

## Acknowledgments

The authors thank Dr. Ernesto T. A. Marques Jr for virus donation, Fabio Jorge Moreira da Silva (ICB/UFRJ) for technical assistance with animal breeding, Julia V. França and Jéssica B. Nascimento-Viana for assistance during data collection.

