## Supplemental Files for "Different outcomes of neonatal and adult Zika virus infection on startle reflex and prepulse inhibition in mice"

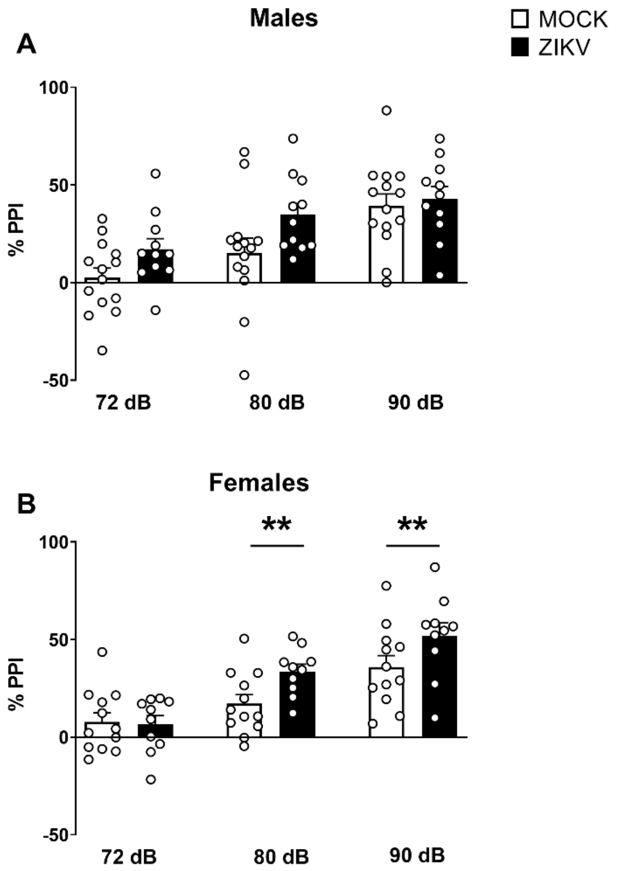


**Supplementary Figure 1. ZIKV infection in the developing brain alters PPI in specific prepulse intensities.** ZIKV did not alter PPI in specific prepulse intensities in males **(A)** but increased PPI in female **(B)** mice in both 80 and 90 dB prepulse intensities (three-way ANOVA, $F_{triple\ interaction(2,86)} = 3.714$, $p = 0.028$). n = 14 Mock and 11 ZIKV for males and 12 Mock and 10 ZIKV for females. **$p < 0.01$ Mock *vs.* ZIKV. Data are expressed as mean ± S.E.M.

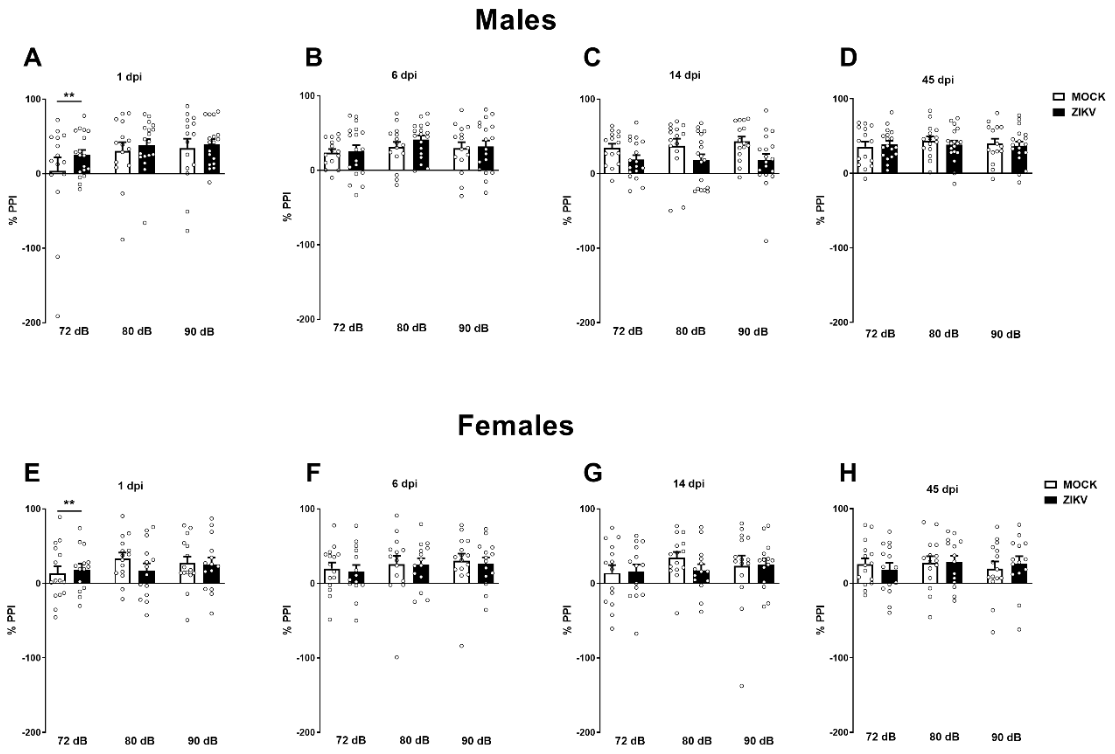


**Supplementary Figure 2. ZIKV infection in the mature brain preserves PPI across time and prepulse intensities.** ZIKV alters PPI of male **(A)** and female adult mice **(E)** one day after infection only at 72 dB prepulse (three-way ANOVA, $F_{\text{pre-pulse x group}(2,116)} = 3.447$, $p = 0.035$). This change is not long-lasting **(B-D and F-H)**. Female mice showed a slightly lower PPI than male at 45 dpi **(D and H)** (three-way ANOVA, $F_{\text{sex}(1,58)} = 4.709$, $p = 0.034$). n = 15 Mock and 18 ZIKV for males and 15 Mock and 14 ZIKV for females. In A: **$p < 0.01$ Mock *vs*. ZIKV. Data are expressed as mean ± S.E.M.

**Table 1S:** Two-way ANOVA results of data from figure 1 (neonatal infection).

| Data | Startle amplitude (Fig. 1A and D) | Startle Latency (Fig. 1B and E) | Average PPI (Fig 1C and F) |
|---|---|---|---|
| Group factor | $F_{(1,43)} = 0.044$<br>$p = 0.834$ | $F_{(1,43)} = 8.149$<br>$p = 0.007$* | $F_{(1,43)} = 5.315$<br>$p = 0.026$ |
| Sex factor | $F_{(1,43)} = 0.345$<br>$p = 0.560$ | $F_{(1,43)} = 2.088$<br>$p = 0.156$ | $F_{(1,43)} = 0.001$<br>$p = 0.975$ |
| Interaction | $F_{(1,43)} = 0.046$<br>$p = 0.831$ | $F_{(1,43)} = 1.046$<br>$p = 0.312$ | $F_{(1,43)} = 0.050$<br>$p = 0.824$ |

**Table 2S:** Three-way ANOVA data results from figure 2 (adult infection).

| Data | Startle amplitude (Fig. 2A and D) | Startle Latency (Fig. 2B and E) | Average PPI (Fig 2C and F) |
|---|---|---|---|
| Group factor | $F_{(1,58)} = 4.864$ *p* = 0.031* | $F_{(1,58)} = 1.591$ *p* = 0.212 | $F_{(1,58)} = 0.146$ *p* = 0.704 |
| Sex factor | $F_{(1,58)} = 4.925$ *p* = 0.030* | $F_{(1,58)} = 0.939$ *p* = 0.337 | $F_{(1,58)} = 2.095$ *p* = 0.153 |
| Dpi factor | $F_{(3,174)} = 3.766$ *p* = 0.012* | $F_{(3,174)} = 2.845$ *p* = 0.039* | $F_{(3,174)} = 0.985$ *p* = 0.401 |
| Group x sex | $F_{(1,58)} = 2.622$ *p* = 0.111 | $F_{(1,58)} = 0.063$ *p* = 0.802 | $F_{(1,58)} = 0.018$ *p* = 0.894 |
| Group x dpi | $F_{(3,174)} = 2.564$ *p* = 0.056 | $F_{(3,174)} = 1.049$ *p* = 0.372 | $F_{(3,174)} = 1.274$ *p* = 0.285 |
| Sex x dpi | $F_{(3,174)} = 1.780$ *p* = 0.153 | $F_{(3,174)} = 2.428$ *p* = 0.067 | $F_{(3,174)} = 0.452$ *p* = 0.716 |
| Triple interaction | $F_{(3,174)} = 2.785$ *p* = 0.042* | $F_{(3,174)} = 0.801$ *p* = 0.495 | $F_{(3,174)} = 1.162$ *p* = 0.326 |

**Table 3S:** Three-way ANOVA results of data from figures 1S and 2S.

| Data | Neonatal data (Fig 1S) | Adult data (Fig 2S) | | | |
|---|---|---|---|---|---|
| | | **1 dpi (Fig. 2SA and E)** | **6 dpi (Fig 2SB and F)** | **14 dpi (Fig 2SC and G)** | **45 dpi (Fig 2SD and H)** |
| Group factor | $F_{(1,43)} = 5.315$ $p = 0.026$* | $F_{(1,58)} = 0.132$ $p = 0.718$ | $F_{(1,58)} = 0.020$ $p = 0.889$ | $F_{(1,58)} = 2.468$ $p = 0.122$ | $F_{(1,58)} = 0.017$ $p = 0.898$ |
| Sex factor | $F_{(1,43)} = 0.001$ $p = 0.975$ | $F_{(1,58)} = 0.436$ $p = 0.512$ | $F_{(1,58)} = 0.713$ $p = 0.402$ | $F_{(1,58)} = 0.666$ $p = 0.418$ | $F_{(1,58)} = 4.709$ $p = 0.034$* |
| Prepulse factor | $F_{(2,86)} = 86.399$ $p < 0.001$* | $F_{(2,116)} = 13.666$ $p < 0.001$* | $F_{(2,116)} = 7.988$ $p = 0.001$* | $F_{(2,116)} = 2.353$ $p = 0.353$ | $F_{(2,116)} = 2.232$ $p = 0.112$ |
| Group x sex | $F_{(1,43)} = 0.050$ $p = 0.824$ | $F_{(1,58)} = 0.726$ $p = 0.398$ | $F_{(1,58)} = 0.255$ $p = 0.615$ | $F_{(1,58)} = 0.956$ $p = 0.331$ | $F_{(1,58)} = 0.015$ $p = 0.902$ |
| Group x prepulse | $F_{(2,86)} = 2.534$ $p = 0.085$ | $F_{(2,116)} = 3.447$ $p = 0.035$* | $F_{(2,116)} = 0.853$ $p = 0.429$ | $F_{(2,116)} = 1.340$ $p = 0.266$ | $F_{(2,116)} = 0.507$ $p = 0.604$ |
| Sex x pre-pulse | $F_{(2,86)} = 0.521$ $p = 0.596$ | $F_{(2,116)} = 1.898$ $p = 0.154$ | $F_{(2,116)} = 1.228$ $p = 0.429$ | $F_{(2,116)} = 1.133$ $p = 0.326$ | $F_{(2,116)} = 0.111$ $p = 0.895$ |
| Triple interaction | $F_{(2,86)} = 3.714$ $p = 0.028$* | $F_{(2,116)} = 0.824$ $p = 0.441$ | $F_{(2,116)} = 0.122$ $p = 0.885$ | $F_{(2,116)} = 1.631$ $p = 0.200$ | $F_{(2,116)} = 2.508$ $p = 0.086$ |
